# Feeling Senseless Sensations: A Crossmodal study of mismatching tactile and virtual visual experience

**DOI:** 10.1101/2024.04.30.591838

**Authors:** Caroline Lehser, Steven A. Hillyard, Daniel J. Strauss

## Abstract

To create highly immersive experiences in virtual reality (VR) it is important to not only include the visual sense but also to involve multimodal sensory input. To achieve optimal results, the temporal and spatial synchronization of these multimodal inputs is critical. It is therefore necessary to find methods to objectively evaluate the synchronization of VR experiences with a continuous tracking of the user. In this study a passive touch experience was incorporated in a visual–tactile VR setup using VR glasses and tactile sensations in mid–air. Inconsistencies of multimodal perception were intentionally integrated into a discrimination task. The participants’ electroencephalogram (EEG) was recorded to obtain neural correlates of visual-tactile mismatch situations. The results showed significant differences in the event-related potentials (ERP) between match and mismatch situations. A biphasic ERP configuration consisting of a positivity at 120 ms and a later negativity at 370 ms was observed following a visual–tactile mismatch. This late negativity could be related to the N400 that is associated with semantic incongruency. These results provide a promising approach towards the objective evaluation of visual–tactile synchronization in virtual experiences.

## 1. Introduction

Virtual reality (VR) is a rapidly growing field with many applications in scenarios such as entertainment, learning and communication. The aim of these applications is to create an immersive experience for the user in an alternative sensory reality. Providing multimodal input can affect the realism and immersion of the experience [1]. A challenge for multimodal VR setups, however, is the integration of multiple devices and the synchronization of multisensory inputs [2].

Some devices have been proposed that would expand a visual VR within a head mounted display (HMD) by including tactile sensory input. Contacting devices like controllers or wearables such as gloves or wristbands that provide vibratory or electrical feedback can be implemented in such VR setups [2, 3].

An alternative to contacted tactile stimulators is the use of contactless tactile feedback. Carter et al. [4] introduced a system that provides multi–point haptic feedback above an interactive surface. This system generates sensations in various shapes in mid–air between 15 cm and 50 cm above the surface using modulated focused ultrasound [5]. Our group (Lehser et al. [6]) demonstrated that somatosensory evoked potentials (SEPs) can be recorded in response to such ultrasound stimuli in mid–air. The recorded SEPs were similar to those elicited by a commonly used contacted vibrating device.

A visual–tactile VR experience with mid–air haptics was developed by Pittera et al. [7] that allowed the user to see and feel rain on their hands. Marchal et al. [8] alsoused mid–air haptics to provide sensations while participants interacted with objects that differed in stiffness in VR. By using virtual haptic sensations in mid–air combined with a visual VR the spatial co–localization with the visual input is very important for immersive experiences. A stimulator that provides contactless feedback is not directly attached to a specific body location, so the tracking of the stimulated body part has to be accurate to achieve a realistic experience.

The evaluation of VR experiences is typically based on the subjective report of the user or on questionnaires. Such assessment during a VR experience is only possible, however, by interrupting the session and breaking the immersion. Thus an objective, non-intrusive evaluation method that would enable the continuous tracking of the user during VR would be highly desirable [1].

One promising approach would be to record electrophysiological measures of violations of the user’s predictions about their interactions with objects in VR. Along these lines, Gehrke et al. [9] demonstrated an ERP-based method for detecting unrealistic VR interactions in visual–tactile sensory integration using the frontal mismatch-negativity (MMN) as an electrophysiological index. Similarly, Singh et al. [10] reported that the realism of hand shape correlates with the MMN. With an increasing realism of the hand shape, the user became more sensitive to subtle inaccuracies in VR. Sensory mismatches can occur in different ways and thus have different effects on perception and neural processing. Unimodal mismatches have been most intensively studied in the auditory modality. In ERP recordings the mismatch negativity (MMN) is typically elicited by deviant sounds (e.g. in frequency or intensity) in a sequence of homogenous stimulations [11, 12]. Analogous MMN responses have been reported in other modalities including vision [13] and touch [14, 15, 16, 17].

A higher order type of mismatch occurs when an event is perceived that is not expected in a particular context. For example, an unexpected or surprising stimulus that is physically different from the expected stimulus may elicit a late positive ERP component usually known as P300. This effect was shown by Kutas et al. [18] in a situation where subjects encountered unexpectedly large letters in words they were reading. In contrast, words that were semantically inappropriate in a sentence context elicited a late negative ERP component called the N400 [18]. The N400 can not only be elicited by the reading of semantically inappropriate words, but also by incongruous spoken and signed words [19]. The N400 can be elicited by a broad class of semantic mismatches in addition to incongruous words; these include anomalies in the form of drawings, photos and videos of faces, objects and actions, as well as sounds and mathematical symbols [19]. In particular, scenes that are semantically incongruent, e.g.,a picture that contains an object that is not expected in the scene, typically elicit an N400 [20]. A similar effect occurs for spatial incongruency, in which the object in the scene is semantically congruent but its location is unexpected [21].

The purpose of the present study was to find an objective electrophysiological index of a multimodal spatial mismatch between haptic and visual stimuli in VR. A visual– tactile VR setup with a head mounted display (HMD) and tactile sensations in mid–air was created for multimodal perception of a touch experience. Spatial inconsistencies of the touch experience were intentionally introduced in a tactile discrimination task. ERPs were recorded to identify neural correlates of visual-tactile mismatch situations in which the mismatch was not relevant to the task.

## 2. Materials and Methods

### 2.1. Participants

A total of twenty subjects participated in this study (mean age: 26.3, 7 female, 13 male). All participants had no known neurological or psychiatric disorder. After a detailed explanation of the procedure, all participants signed a consent form and were told that they could stop the experiment at any time without giving a reason. All measurements were conducted in accordance with the Declaration of Helsinki. The study was approved by the local ethics committee (application: 96/21 Ärztekammer des Saarlandes; Medical Council of the Saarland).

### 2.2. Experimental Setup and Stimuli

Participants were asked to sit relaxed in a chair and to place their hand, the palm facing upwards, on an armrest at the table in front of them. The index, middle and ring fingers of the right hand were placed at marked positions. To avoid movement of the fingers out of the focus of the tactile stimulator the finger positions were demarcated by partitions. The setup is shown in figure 1 (left). The tactile stimulations were generated by an ultrasonic board (Ultraleap Stratos Explore, Ultraleap Ltd., England) introduced by Carter et al. [4]. The ultrasonic board consisted of an array of 16x16 ultrasonic transducers, which generated haptic sensations in mid–air with modulated focused ultrasound. For the tactile stimuli in this study, a line with a length of 2 cm was generated using time point streaming and a modulation frequency of 200 Hz. The board was placed 17 cm above the participant’s right hand. The location of the stimuli was adjusted to the positions of the participants index, ring or middle finger so that each stimulation could only be perceived on the intended finger.

**Figure 1.**
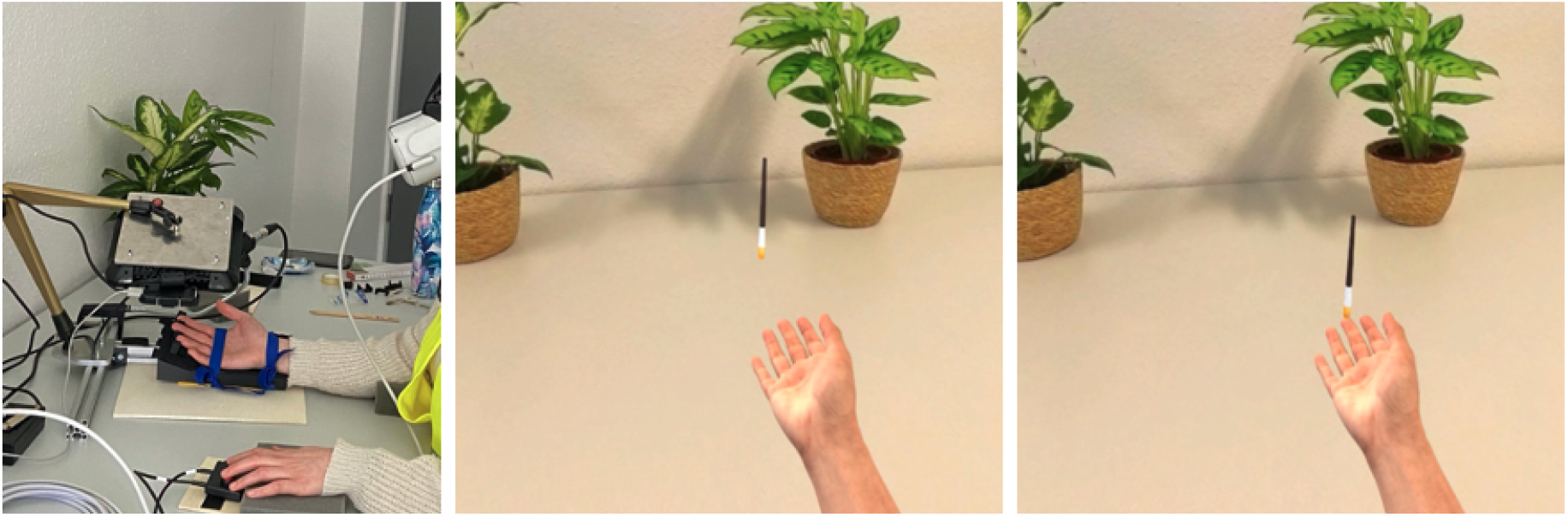
Left: General setup: Participant wearing VR glasses, headphones and the wireless EEG–cap, with the right hand at rest under the stimulator and the left hand responding by pressing the buttons; Middle: View seen by the participant in VR with the virtual hand and the small paintbrush in resting position; Right: The paintbrush moves and touches a finger.

Participants wore VR glasses (Meta Quest 2, Meta) for the visualization of the task situation. The environment in virtual reality was built with a 360° picture of the physical room where the study took place to enhance the immersive experience. On the table in VR the participants saw a hand and forearm. The position of the virtual hand was set individually for each participant so that the virtual arm and hand matched the position of the real arm. The movement of the real fingers and hand was tracked by a LeapMotion (Ultraleap Ltd., England) and transmitted to the virtual hand. A small virtual paintbrush was positioned above the virtual hand. On each trial the virtual brush moved to a finger and touched it in synchrony with the real tactile sensation. Figure 1 (middle) shows the position of the hand and the brush during the inter–stimulus interval. A trial began when the brush began to move towards a finger; this movement lasted 120 ms until the moment of touch. Figure 1 (right) shows the moment of the brush touching the finger. The brush touched the finger for 200ms, which was the same duration as the mid–air tactile stimulation.

The experiment was divided into six blocks of 210 trials each, which resulted in a total of 1260 stimulations. The probability of a stimulation was 33,3 % for each of the three fingers. The participant’s task was to decide on which finger the tactile sensation was felt and to push with the left hand one button for the sensation at the index finger and a second button for sensations at either the middle or ring finger. On most of the trials the virtual brush touched the same finger that was tactilely stimulated. But for 20% of the index finger stimulations a mismatch between the tactile stimulation and the visual virtual reality was produced. For these mismatch trials the tactile stimulation was felt on the index finger but the brush was seen to touch the middle finger. This mismatch was not task relevant, so that the associated ERP could more specifically reflect the modality mismatch rather than a task-related decision.

To avoid any interference from acoustic noise, especially the sound generated by the tactile stimulator, the participants wore earplugs as well as headphones that presented a pink masking noise at a soft and comfortable level. Participants could take breaks between the measurement blocks as they wished to avoid fatigue and to move their right hand.

### 2.3. Data Acquisition and Analysis

ERPs were recorded with a commercially available 64–channel wireless EEG cap with reference to the right earlobe (g.tec g.nautilus, Guger Technologies Austria) using a sampling frequency of 500 Hz. Electrode impedances were below 30 kΩ. Physiological data was analyzed using MATLAB 2023b (The MathWorks, Inc). A zero–phase FIR bandpass filter of order 1000 and cut–off frequencies of 1 and 30 Hz was applied to the raw EEG signal. ERPs were trial–wise baseline corrected by subtracting the mean potential between 200 ms and 150 ms before the onset of the tactile stimulation. Noisy channels were interpolated by visual inspection based on their time course and power spectra. These channels were interpolated using EEGLAB’s spherical interpolation method (version 2023.0 [22]). Trials with artefacts were removed using an amplitude threshold of ± 60 *μ*V. The data from the six measurement blocks were pooled together and sorted according to the four stimulation conditions:three matching conditions (stimulation to the index, middle or ring finger) and one mismatch condition (tactile stimulus to index finger with visual brush touching middle finger).

### 2.4. Statistical Analysis

Statistical analyses comparing ERP amplitudes on the different matching trials with the mismatch trials within participants were performed with MATLAB using two–tailed t– tests. The significance threshold for the t–tests was set to *α <* .05. *p–value* adjustments for multiple comparisons were made following the Benjamin–Hochberg FDR correction procedure [23]. Reaction times between the different conditions (matching trials in different positions and mismatching trials) were also statistically analyzed with two– tailed t–tests (*α <* .05).

## 3. Results

The grand average ERPs over all participants were calculated over the following numbers of artefact free trials: 70 mismatch trials, 260 index–match trials, 300 middle–match trials and 300 ring–match trials. Figure 2 shows the grand average waveforms at electrode Cp3 for all conditions. Electrode Cp3 was chosen to illustrate the effects of mismatching tactile stimuli because it overlies the somatosensory cortex contralateral to the stimulation.

**Figure 2.**
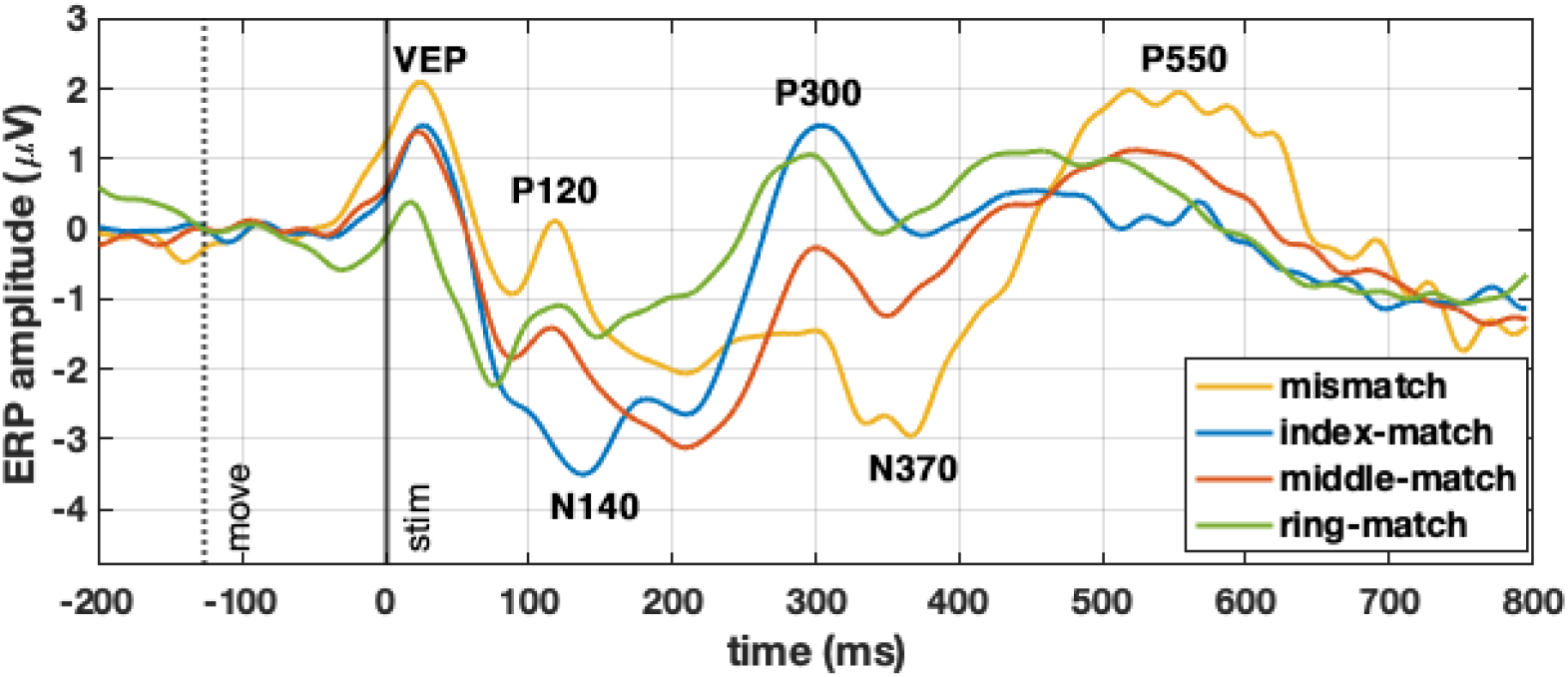
Grand averages of the ERP waveforms for electrode Cp3 for mismatch (yellow), index–match (blue), middle–match (red), ring–match (green) stimulation with labels of the peaks. The vertical dotted line indicates the start of the movement of the virtual brush, the vertical solid line indicates the onset of the tactile stimulation at the same time the brush touches the finger in VR.

The dotted vertical line indicates the start of the movement of the brush in the VR visualization while the solid vertical line indicates the time point when the tactile stimulation began and the brush touched the finger in VR. Around 30ms after this time point a positive peak was present in all conditions, which represents the visual evoked potential (VEP) elicited by the movement of the brush. This positivity was followed be a smaller positive peak at around 120 ms (P120) under all conditions except for the index–match, which showed a negative wave at 140ms (N140). A subsequent positive wave at 300 ms (P300) was elicited by all three of the matches. In contrast, the mismatch showed a distinct negative wave with a maximum amplitude at 370 ms (N370) followed by a broad positive wave (P550).

Figure 3 (top) compares the ERPs elicited by index–match and mismatch recorded at electrode Cp3. The gray areas indicate the time ranges when the difference between the match and mismatch ERP averages were significant (*p <* 0.05). Figure 3 (bottom) shows the difference wave formed by subtracting mean of the ERP to the index–match from the ERP to the mismatch. The most prominent components were a significant positive difference between 70 ms and 160 ms followed by a significant negative difference from 240 ms to 470 ms. A very late positivity between 500 and 650 ms was also significant.

**Figure 3.**
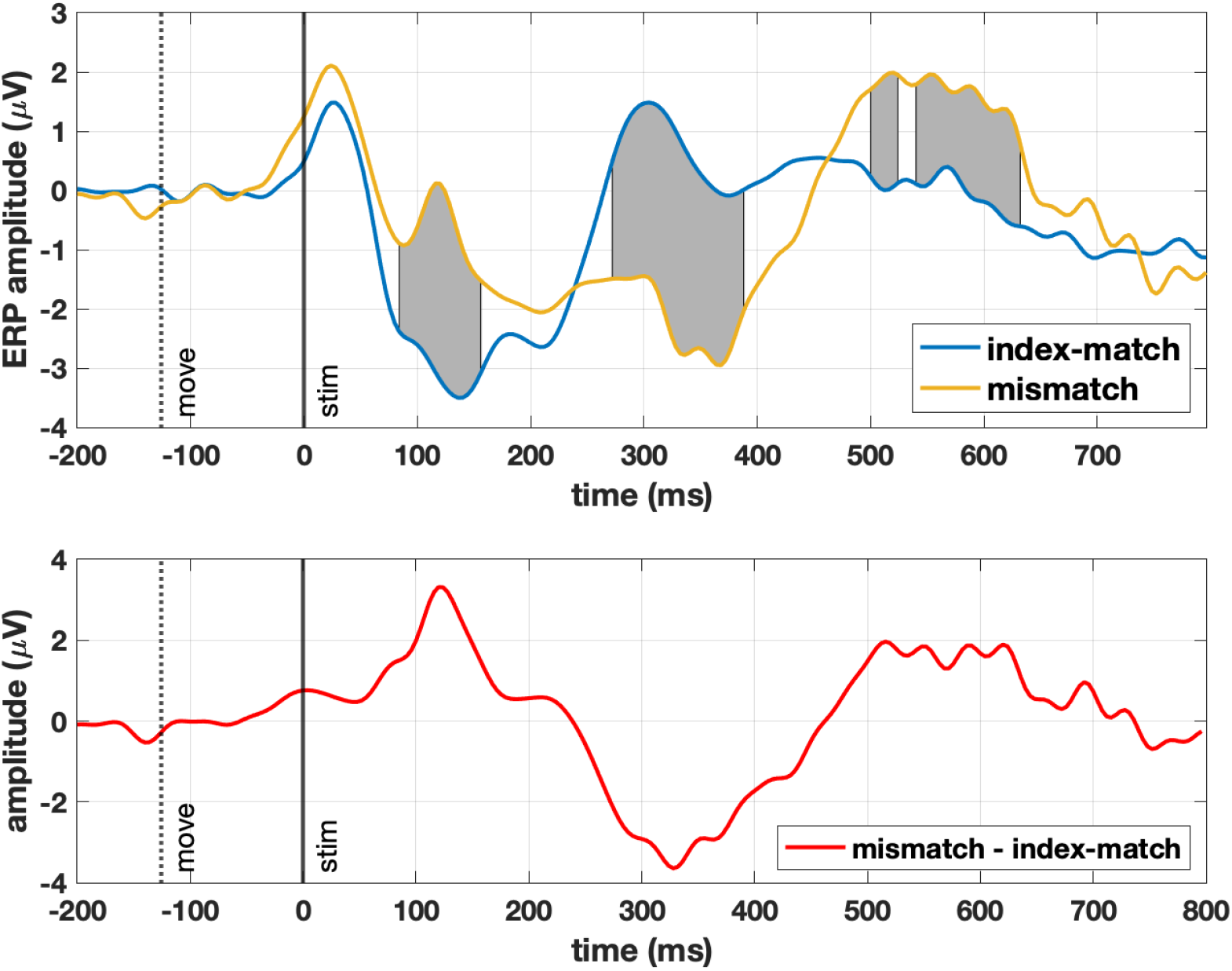
Top: Grand averages of the ERP waveforms for electrode Cp3 for mismatch (yellow), index–match (blue). The gray areas indicate the time range with significant differences (*p <* 0.05) between the index–match and the mismatch. The vertical dotted line indicates the start of the movement of the virtual brush, the vertical solid line indicates the onset of the tactile stimulation at the same time the brush touches the finger in VR. Bottom: The difference wave formed by subtracting the average ERP for index–match from the average ERP for the mismatch.

In figure 4 topographic plots are shown with 64 channels averaged over 50 ms intervals at the latencies of the critical mismatch effects. In the 100–150 ms interval the index–match showed a centro-parietal negativity (N140) contralateral to the stimulated finger that is characteristic of the somatosensory ERP [24]. In sharp contrast, the mismatch elicited a large, centrally distributed positivity (P120) that was also present to a lesser extent to the middle–match. The 300–350 ms interval showed a posteriorly distributed positivity (P300) for all matching stimuli. In contrast, the mismatch elicited a large anteriorly distributed negativity. This negativity (N370) associated with the mismatch became larger and more widespread in scalp distribution in the 350–400 ms interval. The late positivity (P550) associated with the mismatch had a widespread anterior scalp distribution.

**Figure 4.**
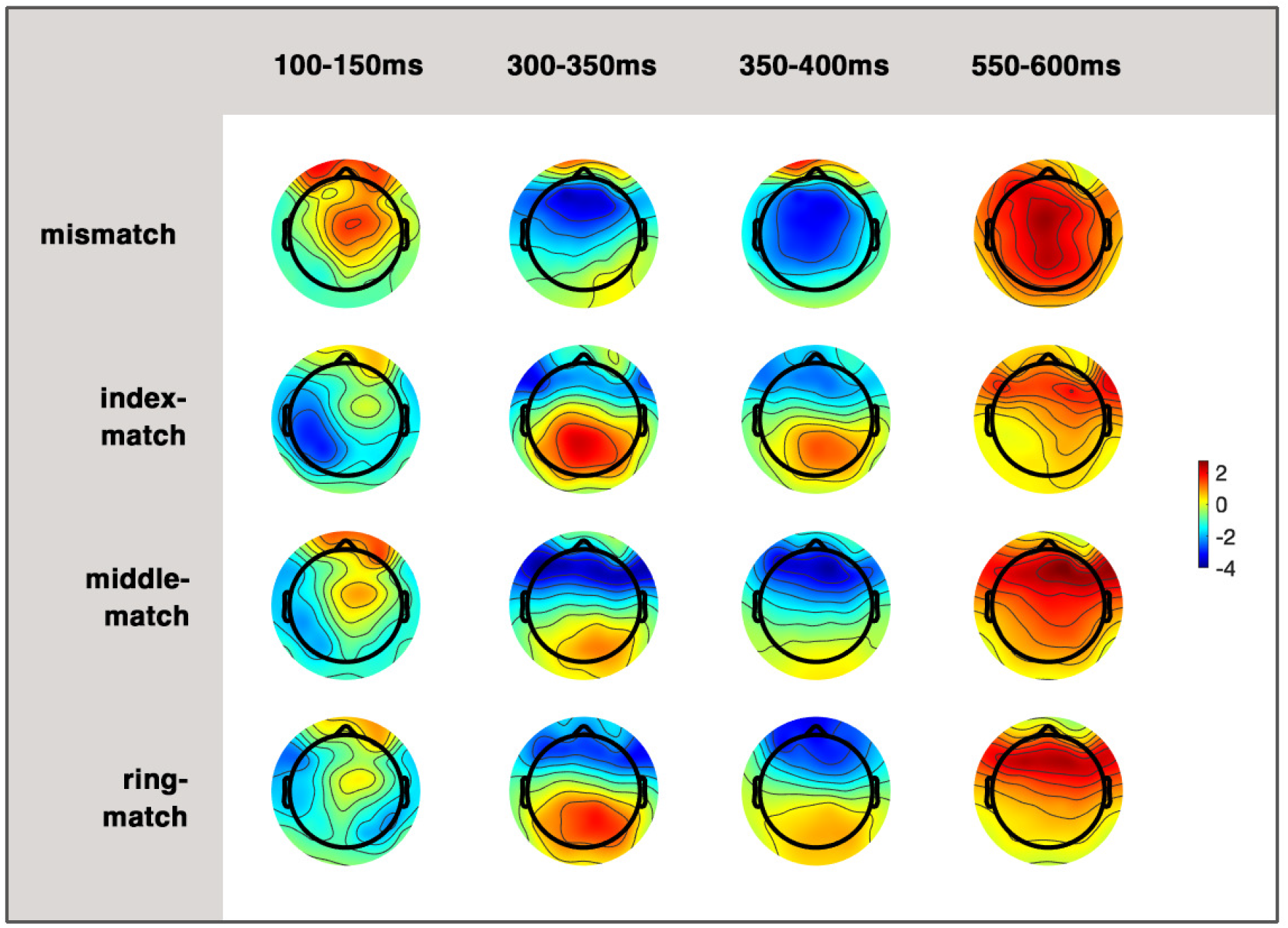
Topographic plots of 64 channels for the ERP averaged over 50ms intervals. Each row shows the topographic plots of one condition over four different time ranges. The averaged amplitude values in *μV* are color-coded according to the color bar on the right.

In figure 5 topographic plots are shown with 64 channels averaged over 50 ms intervals for the difference ERP between the index–match and the mismatch conditions for four different time ranges. The bottom row shows plots of statistical significance, where red dots indicate a significant difference (*p <* 0.05) for the corresponding electrode. The 100–150 ms interval showed a widespread positivity, which was maximal in the central parietal region, but significant for almost all electrodes. The 300–350 ms and 350–400 ms intervals showed a widespread negativity with a centro–parietal maximum that was significant over the entire posterior scalp. The 550–600 ms interval showed a widespread positivity, that is significant in the central parietal region.

**Figure 5.**
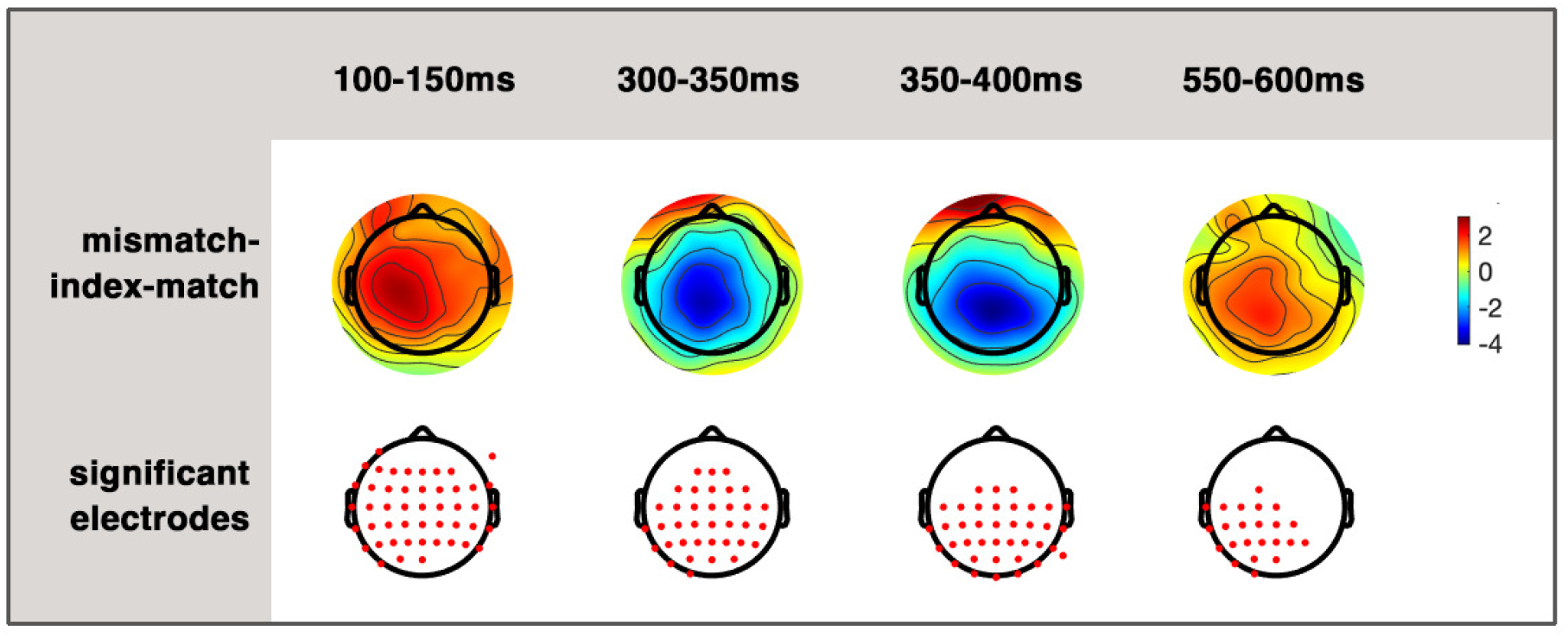
Topographic plots of the difference ERP between index–match and mismatch conditions averaged over 50ms intervals and four different time ranges.Bottom row: Significance plots of the difference between index–match and mismatch. Red dots indicate a significant difference for the corresponding electrode (*p <* 0.05).

Figure 6 shows the reaction times (RTs) (i.e. the time between the tactile stimulation and the button response) under the different conditions. The match trials had mean RTs between 445 ms and 517 ms, while the mismatch trials had a mean RT of 688 ms. The mismatch RT was significantly slower than the middle–match RT (*p <* 10^*–*6^) and both the index–match and ring–match RTs (*p <* 10^*–*8^). The middle-match RT was significantly slower than both the index–match and ring–match RTs (*p <* 10^*–*8^), which did not differ from each other.

**Figure 6.**
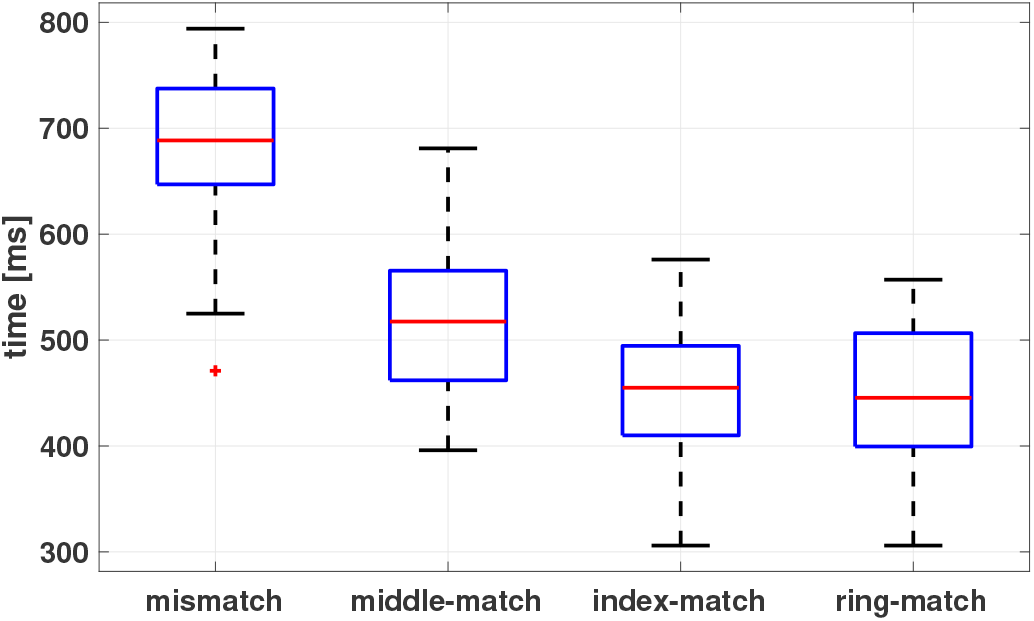
Mean reaction times of button presses in response to the tactile stimulation for each condition. The median, maximum and minimum, and the first and third quartiles of the reaction time are illustrated.

## 4. Discussion

The aim of the present study was to work towards an objective, non-invasive evaluation method for a spatial visual–tactile mismatch in a multimodal VR experience. In particular, ERP recordings were used to index a mismatch between tactile stimulation provided by a contactless stimulator and the visualization of this stimulation in visual VR during a tactile discrimination task. The participant’s task was simply to judge the position of a tactile stimulus and the question of whether the visual and tactile stimuli matched in localization was not task relevant. The results showed that the ERP to the mismatch was strikingly different from the ERPs to the matching multimodal stimulation.

The early positive wave labeled as VEP in figure 2 was a component of the visual evoked potential elicited by the movement of the brush shown in visual VR that was equivalent for all conditions. This movement started 120 ms before the tactile stimulation and thus elicited an earlier response compared to the tactile stimulation for all conditions.

The difference wave for the ERPs of the index–match subtracted from the ERPs of the mismatch showed two main time ranges with large differences that were statistically significant over widespread scalp areas. The earliest neural sign of the modality mismatch was a sharp positivity peaking at 120 ms (P120). Since the index–match and mismatch, had the same tactile stimulation at the same location and only differed in the visual stimulation it can be assumed that the ERP difference is a consequence of the sensory mismatch. The ERP to the index–match showed a contralateral N140, which is typically recorded after tactile stimulation. This wave was overridden by the P120 triggered by the sensory mismatch. The P120 appears similar to the positive mismatch component reported by Strömmer et al. [17], who recorded ERPs in response to the tactile stimulation of two fingers in an oddball paradigm. Stimulation of the infrequently stimulated finger elicited a positive somatosensory mismatch component over 100–300 ms with a centro–parietal distribution. Shinozaki et al. [16] also recorded a change–related positivity over 100–200 ms during a similar paradigm. Thus, it appears that an early positivity starting at around 100 ms may be a general characteristic of the ERP response to a mismatch in the somatosensory modality.

Following the P120, the mismatching condition elicited a late negativity (N370) over the centro–parietal scalp, This negativity contrasted sharply with the ERP to the match conditions, which showed a P300 wave during this time interval. In sensory decision tasks the task–relevant stimuli typically elicit the P300 component. The present design, however, included a visual mismatching stimulus that was irrelevant to the task of discriminating the position of the tactile stimulus. This visual-tactile mismatch converted the typical P300 into a late negativity (N370), which could be related to the N400 that has been observed after semantically incongruent stimulation.. As reviewed in the Introduction, there have been a wide range of studies that have reported an N400 [19]. The key characteristic of situations that elicit an N400 appears to be a violation of a semantic context, either in language or in real-world situations. To our knowledge, however, there have been no studies that analyzed the N400 during a sensory mismatch in a multimodal VR scenario. The present results suggest that incongruent sensory stimulation in different modalities (i.e., senseless sensations) may be included in the category of real-world violations that elicit an N400. Further studies are needed to determine the range of situations where incongruent stimulation produces an N400.

When the brush started moving towards a finger, there was a point in time when it could be visually determined which finger was going to be touched by the brush. The middle finger was visually touched by the brush in two situations, the middle–match and the mismatch (when the index finger received the tactile touch). So when the brush was seen approaching the middle finger there was uncertainty as to which finger would receive the tactile stimulus. This uncertainty of the visual middle finger that was resolved by the tactile stimulation may be responsible for eliciting the small negativity (N370) when it turned out to be a match and the much larger negativity when it resulted in a mismatch.

This uncertainty explanation was supported by the reaction time data., The mean reaction times were shortest and not significantly different for the index–match and the ring–match conditions. In both cases there was complete certainty that the visual and tactile stimuli would match. The mean reaction time to the mismatch was the slowest of all, due to the need for processing the mismatching sensory information. But the mean reaction time to the middle–match was significantly longer than the reaction time to the other matches, which was likely a consequence of the above mentioned uncertainty.

## 5. Conclusion

In this study an environment of visual and tactile VR was designed to objectively evaluate a spatial mismatch in a multimodal VR experience. The visual VR of an HMD was coordinated with an ultrasound based tactile stimulator. During the VR experience, situations occurred where the sensory input of the visual and tactile modality did not match. Prominent ERP correlates of these mismatch situations were observed. An early positivity (P120) and a later negativity (N370) were found to be significantly different from the matching situations. These components appeared to be robust indicators of the lack of spatial congruency of the visual and tactile stimulation. The experimental setup used in the present study enables a wide range of possibilities for generating mismatching situations in tactile and visual VR. The effects of the synchronization of different modalities in VR can be analyzed not only regarding the location of the stimulation but also the temporal correspondence of multi-modal events.

## Acknowledgments

This work was partially supported by the German Federal Ministry of Education and Research, Grant 13FH050KX1 and by the European Union [Europaeischer Fonds fuer regionale Entwicklung (EFRE)]

